# Tissue-specific and endogenous protein labeling with split fluorescent proteins

**DOI:** 10.1101/2024.02.28.581822

**Authors:** Gloria D. Ligunas, German Paniagua, Jesselynn LaBelle, Adela Ramos-Martinez, Kyle Shen, Emma H. Gerlt, Kaddy Aguilar, Alice Nguyen, Stefan C. Materna, Stephanie Woo

## Abstract

The ability to label proteins by fusion with genetically encoded fluorescent proteins is a powerful tool for understanding dynamic biological processes. However, current approaches for expressing fluorescent protein fusions possess drawbacks, especially at the whole organism level. Expression by transgenesis risks potential overexpression artifacts while fluorescent protein insertion at endogenous loci is technically difficult and, more importantly, does not allow for tissue-specific study of broadly expressed proteins. To overcome these limitations, we have adopted the split fluorescent protein system mNeonGreen2_1-10/11_ (split-mNG2) to achieve tissue-specific and endogenous protein labeling in zebrafish. In our approach, mNG2_1-10_ is expressed under a tissue-specific promoter using standard transgenesis while mNG2_11_ is inserted into protein-coding genes of interest using CRISPR/Cas-directed gene editing. Each mNG2 fragment on its own is not fluorescent, but when co-expressed the fragments self-assemble into a fluorescent complex. Here, we report successful use of split-mNG2 to achieve differential labeling of the cytoskeleton genes *tubb4b* and *krt8* in various tissues. We also demonstrate that by anchoring the mNG2_1-10_ component to specific cellular compartments, the split-mNG2 system can be used to manipulate protein function. Our approach should be broadly useful for a wide range of applications.

## Introduction

Protein labeling by fusion with genetically encoded fluorescent proteins has been a powerful tool for studying biological processes, allowing scientists to visualize and track proteins of interest in live cells. Fluorescent protein labeling has been especially useful for investigating the dynamic processes that occur during embryonic development. However, traditional methods for generating and expressing fluorescent fusion proteins, especially in multicellular organisms, have several drawbacks. In zebrafish and other model organisms, expression of fusion proteins can be achieved by injection of *in vitro* transcribed mRNA (Rosen et al., 2009), which is ubiquitous, or by transgenesis, which utilizes gene regulatory elements to drive spatiotemporal restricted expression (Clark et al., 2011). These approaches, however, run the risk of producing overexpression artifacts, in which proteins may not function or localize correctly when expressed at higher than wild-type levels (Simiczyjew et al., 2014). An alternative approach is to knock in fluorescent protein coding sequences into the genetic locus of that protein of interest (Albadri et al., 2017; Auer and Del Bene, 2014; Kimura et al., 2014). Although this approach has the advantage of preserving endogenous regulation of that protein’s expression, many proteins are expressed broadly; issues arise when there is a need to study a broadly expressed protein in a specific tissue. Thus, there is a need for tissue-specific and endogenous tagging of proteins.

Split fluorescent proteins (split-FPs) are self-complementing protein fragments that only fluoresce when bound together. Split-FPs have been successfully used to visualize and quantify cell-cell interactions (Feinberg et al., 2008), signaling pathway activation (Harvey and Smith, 2009), and subcellular protein localization (Cho et al., 2022). One commonly used split-FP system is based on the yellow-green fluorescent protein monomeric NeonGreen2 (mNG2) in which strands 1–10 of the mNG2 beta-barrel (mNG2_1-10_) and strand 11 (mNG2_11_) are expressed as independent protein fragments (Feng et al., 2017). On their own, the fragments are nonfluorescent, but when present in the same cell, they will self-assemble into a bimolecular complex with similar spectral properties to the intact, full-length fluorescent protein. The split-mNG2 system has been demonstrated to function in several different organisms and cell types (Cho et al., 2022; Kesavan et al., 2021; O’Hagan et al., 2021). Here, we adapt it for use in zebrafish to achieve tissue-specific and endogenous protein labeling. In our approach, mNG2_1-10_ is expressed under the control of a tissue-specific promoter using standard zebrafish transgenesis techniques. Because the mNG2_11_ fragment is only 16 amino acids long, its short sequence can be easily inserted into endogenous genetic loci by CRISPR/Cas-directed gene editing. In this way, the mNG2_11_-tagged protein will continue to be expressed at endogenous levels, but fluorescent signal will only be detected in tissues in which mNG2_1-10_ is co-expressed (Fig. 1A).

**Figure 1.**
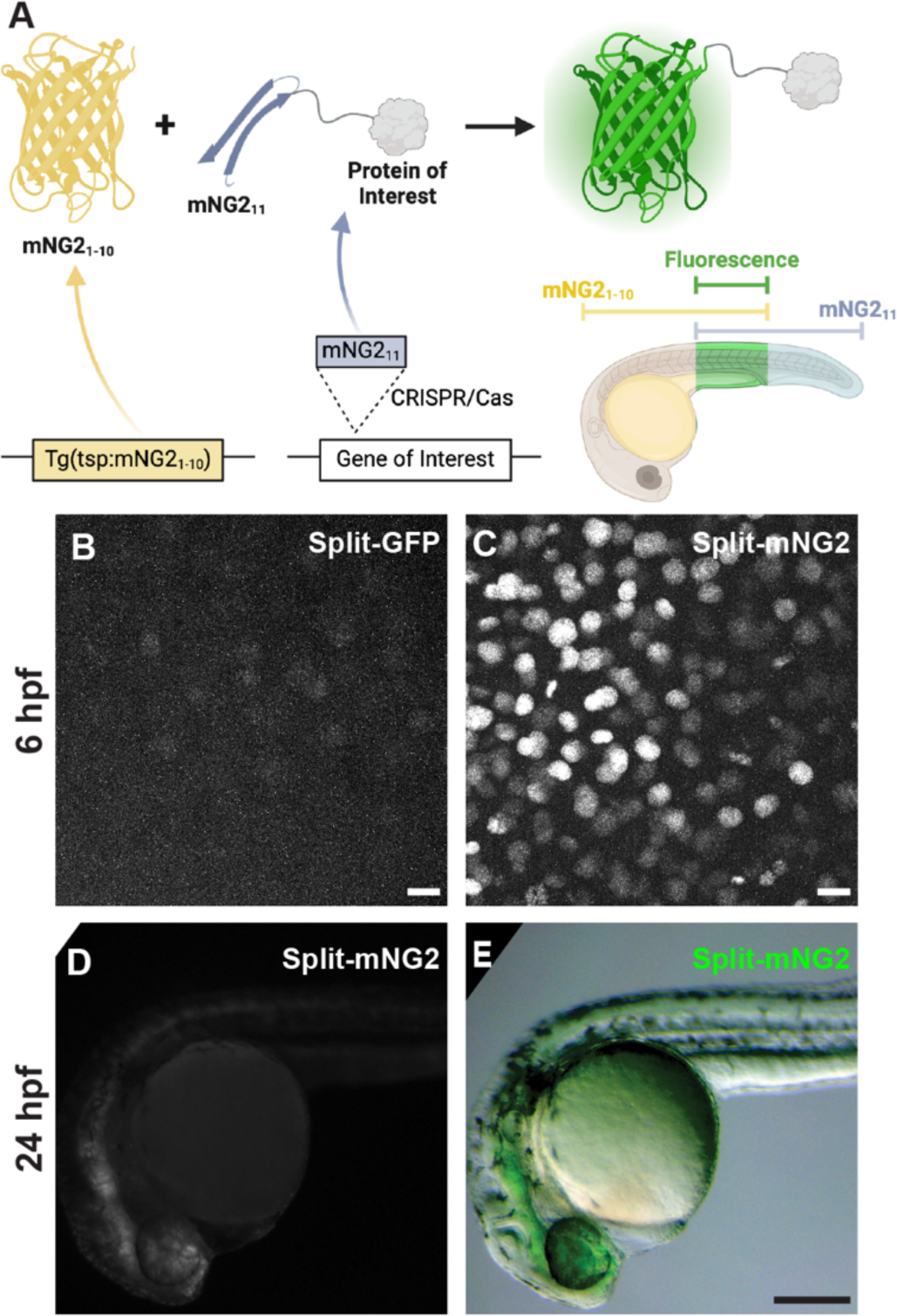
Split fluorescent protein fragments are functional in zebrafish embryos. **A.** Schematic illustrating our protein labeling strategy using a split fluorescent protein. Transgenic (Tg) mNG21-10 is expressed under the control of a tissue-specific promoter (tsp) while mNG211 is inserted into protein-coding genes by CRISPR/Cas-directed gene editing. Fluorescence (green) is only generated in tissues co-expressing mNG21-10 and the mNG211-tagged protein of interest. **B–E.** Embryos were injected with GFP1-10 and GFP11-H2B (split-GFP, B) or mNG21-10 and mNG211-H2B (split-mNG2, C–E) mRNAs then imaged at 6 hours post-fertilization (hpf) on a confocal microscope (B, C) or at 24 hpf on a fluorescence stereomicroscope (D, E). Scale bars in B and C, 50 µm. Scale bar in E, 200 µm.

## Results

### mNG2_1-10_ and mNG2_11_ can assemble fluorescent complexes in zebrafish embryos

To assess the viability of our protein labeling strategy, we first determined if split-FP fragments could self-assemble in zebrafish embryos to form functional fluorescent complexes (Fig. 1B–D). We tested two different FP_1-10/11_-type systems, split-GFP (Kamiyama et al., 2016) and split-mNG2 (Feng et al., 2017). We injected mRNAs encoding GFP_1-10_ and GFP_11_-H2B (GFP_11_ fused to histone 2B) or mNG2_1-10_ and mNG2_11_-H2B (mNG2_11_ fused to histone 2B) into zebrafish embryos. For both systems, expression of the FP_1-10_ or FP_11_ fragments alone did not produce fluorescence. However, when both fragments were co-expressed, we could detect nuclear-localized fluorescent signals by 6 hours post-fertilization (hpf) using confocal fluorescence microscopy (Fig. 1B–C). We observed that embryos expressing split-mNG2 fragments (Fig. 1C) were brighter than those expressing the split-GFP fragments (Fig. 1B), consistent with previous studies showing that split-mNG2 exhibits improved signal-to-background ratios compared to split-GFP (Feng et al., 2017). This difference in brightness persisted over time so that by 24 hpf split-mNG2 fluorescence was bright enough to be detected by a fluorescence stereomicroscope (Fig. 1D–E). Based on these observations, we only used the split-mNG2 system for further experiments.

### Generating mNG2_1-10_ transgenic lines

We next determined whether transgene-driven expression of mNG2_1-10_ could be used to spatially restrict fluorescence (Fig. 2). We generated multiple transgenic zebrafish lines that express mNG2_1-10_ under control of various promoters representing a broad range of tissue types including *fezf2* (brain and eye) (Berberoglu et al., 2009), *myl7* (myocardium) (Huang et al., 2003), and *ubb* (ubiquitous expression) (Mosimann et al., 2011). To verify that these transgenic lines were functional, we injected transgenic embryos with mNG2_11_-H2B mRNA, which would be distributed ubiquitously, and qualitatively assessed fluorescence patterns at 24 or 48 hpf. We found that uninjected mNG2_1-10_ transgenic embryos exhibited no detectable fluorescence (Supplemental Fig. 1). In contrast, transgenic embryos injected with mNG2_11_-H2B mRNA exhibited fluorescence in spatially restricted patterns consistent with the promoter used to drive mNG2_1-10_ expression (Fig. 2A–F). Moreover, these fluorescence patterns were qualitatively similar to those in embryos expressing full-length, intact GFP under control of the same tissue-specific promoters (Fig. 2G–L).

**Figure 2.**
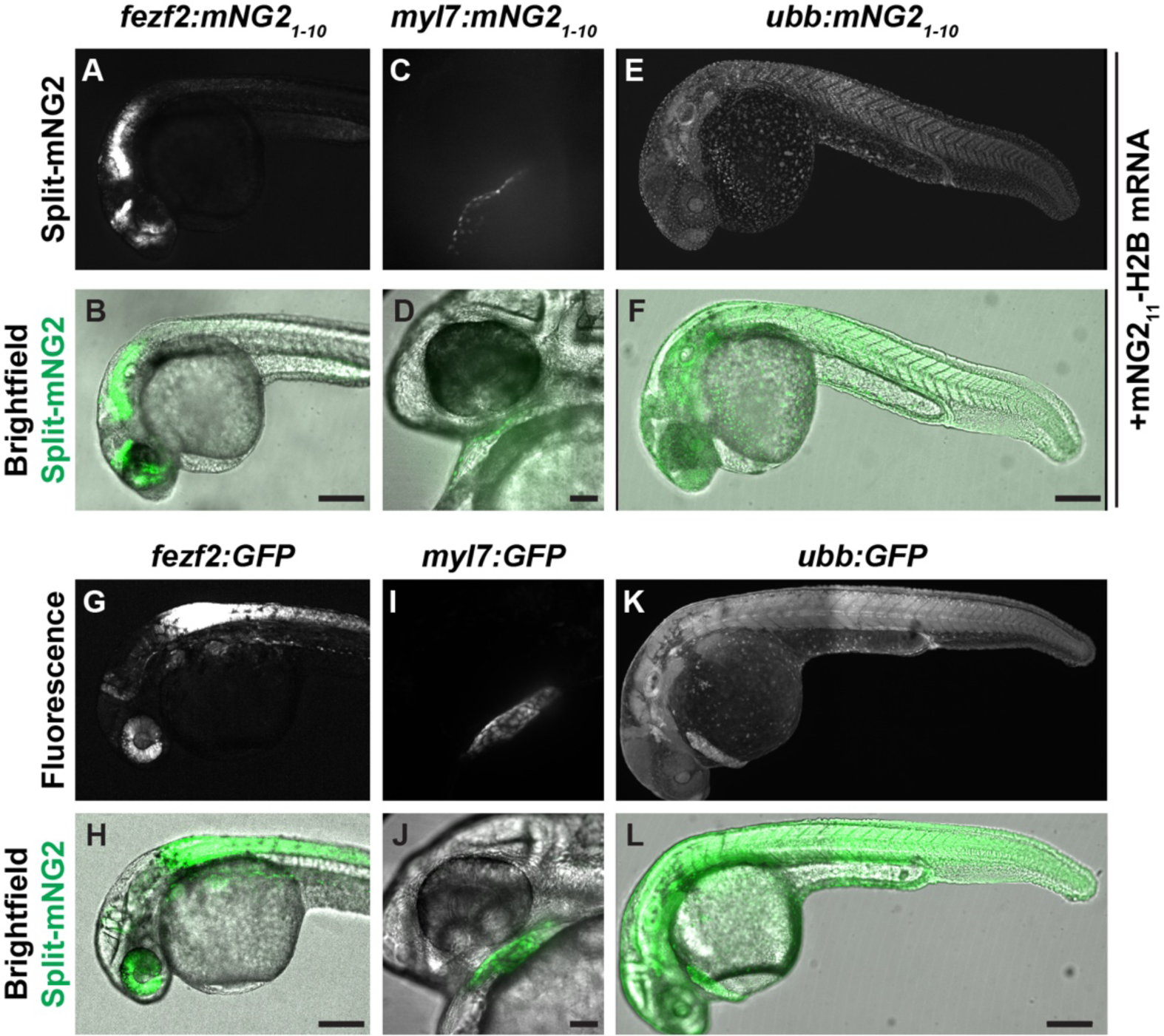
Split fluorescent protein labeling can be spatially restricted by transgenic expression of mNG21-10. **A–F.** Transgenic embryos expressing mNG21-10 under control of the *fezf2* (A–B), *myl7* (C–D), or *ubb* (E–F) promoters and injected with mNG211-H2B mRNA. **D–F.** Transgenic embryos expressing GFP under control of the *fez1* (G–H), *myl7* (I–J), or *ubb* (K–L) promoters. Images were acquired at 24 hours post-fertilization. Scale bars in B, F, H, and L, 200 µm. Scale bars in D, J, 50 µm.

### mNG2_11_ tagging by CRISPR/Cas-directed gene editing

We next determined whether proteins of interest could be tagged with mNG2_11_ at their endogenous genetic loci by CRISPR/Cas-guided homology directed repair (Fig. 3). Previous reports have suggested that split-FP tagging works best for highly expressed genes (Goudeau et al., 2021; O’Hagan et al., 2021). Therefore, we targeted three genes that are highly expressed with relatively broad patterns — *tubb4b*, which codes for Beta-tubulin 4b; *krt8*, which codes for Keratin 8; and *h2az2b*, which codes for histone H2A. We designed guide RNAs (gRNAs) targeting each gene just downstream of the start (*tubb4b*) or upstreak of the stop (*krt8, h2aza2b*) codon to generate, respectively, N- or C-terminal mNG2_11_ tags. We injected gRNAs together with Cas9 mRNA and a repair template that contained the coding sequence for mNG2_11_ and a short linker (Fig. 3A); the repair template consisted of double-stranded DNA with single-stranded homology arms of p30 bp at each end (Liang et al., 2017). To verify that the knock-in was successful, we pooled injected embryos and performed insert-specific PCR that amplified the mNG2_11_ insertion but not the unedited wild-type (Fig. 3B).

**Figure 3.**
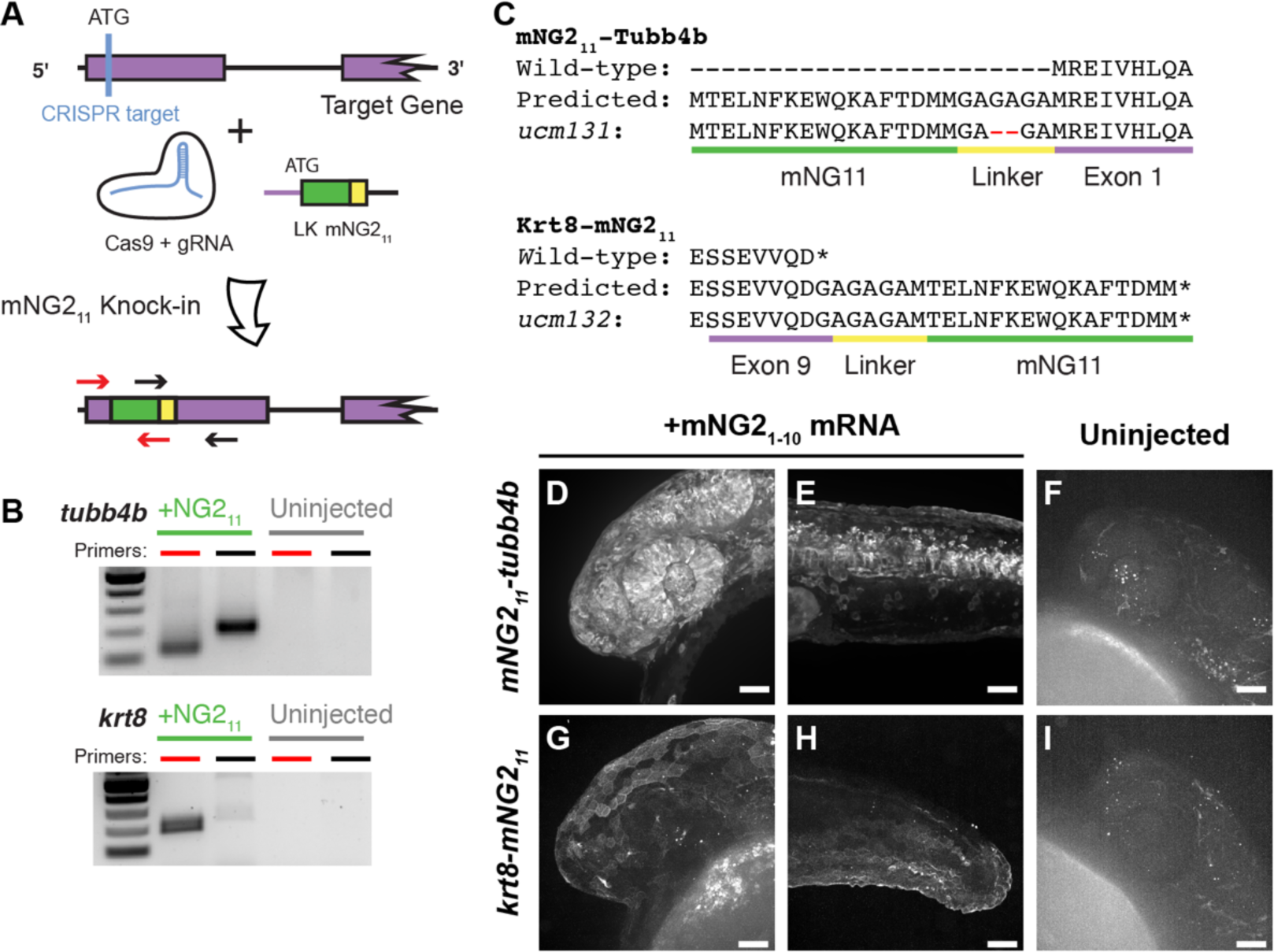
mNG211 tagging by CRISPR/Cas-directed gene editing. **A.** Schematic of CRISPR/Cas-directed mNG211 insertion into target genes. Purple, endogenous exon sequence. Yellow, linker (LK). Green, mNG211. ATG, start codon. Arrows denote primers used in B. **B.** mNG211 insertion was assessed by PCR. The primers used correspond to the arrows shown in A. **C.** Amino acid sequences of wild-type, predicted mNG211 fusions, and recovered alleles for Tubb4b and Krt8. Mismatches between the predicted and recovered sequences are highlighted in red. Asterisks, stop codons. **D–I.** Representative images of *mNG211-tubb4b* (D-F) and *krt8-mNG211* (G–I) embryos injected with mNG21-10 mRNA (D–E, G–H) or uninjected (F, I). Images were acquired at 24 hours post-fertilization. Images in F and I have been overexposed to emphasize lack of fluorescence. Scale bars, 50 µm.

For *tubb4b*, we estimated the knock-in efficiency using quantitative PCR. To determine mNG2_11_ prevalence, we pooled and extracted DNA from 30 injected F_0_ embryos at 24 hpf. We amplified mNG2_11_ using insert-specific primers and amplified the untargeted, single-copy gene *prox1a* for comparison; we obtained a ΔCt of 5 cycles between the two. As zebrafish are diploid, *prox1a* is present in two copies per cell, but each mNG2_11_ knock-in likely occurred only in one *tubb4b* allele per cell. We thus estimated that roughly 1 in every 16 cells in our pooled sample carried the knock-in allele, corresponding to a knock-in efficiency of about 6%, although not necessarily in-frame nor equally distributed among embryos. This knock-in efficiency is on par with other reports of CRISPR-guided knock-in in zebrafish (Auer and Del Bene, 2014; Zhang et al., 2023).

To establish stable, germline-transmitted lines for the mNG2_11_ insertions, we raised injected F_0_ fish to adulthood and identified several founders representing multiple alleles for each gene. Some alleles contained indel mutations at the insertion junctions or within the insertion itself. For example, both alleles recovered for *h2az2b* contained mutations within the mNG2_11_ sequence and produced very dim fluorescence (Supplemental Fig. 2). Therefore, we chose to propagate only alleles with precise integration of the mNG2_11_ sequence, resulting in establishment of one line each for *tubb4b* (*tubb4b^ucm131^*, referred to here as *mNG2_11_-tubb4b*) and *krt8* (*krt8^ucm132^*, referred to here as *krt8-mNG2_11_*).

To confirm that the mNG2_11_ tag is functional and does not alter endogenous expression patterns, we injected embryos with mNG2_1-10_ mRNA and qualitatively assessed fluorescence. For *mNG2_11_-tubb4b*, we observed strong fluorescence at 24 hpf that was especially prominent in the eye and brain (Fig. 3D) and along the neural tube (Fig. 3E). For *krt8-mNG2_11_*, fluorescence appeared restricted to the skin epidermis at 24 hpf (Fig. 3G, H). These fluorescence patterns are consistent with the reported expression patterns for both *tubb4b* (Thisse and Thisse, 2008; Zhuo et al., 2012) and *krt8* (Fischer et al., 2014; Thisse and Thisse, 2008). At the subcellular level, we observed that fluorescence for both genes was enriched at the cell periphery and excluded from the nucleus, which would be expected for cytoskeletal filaments. For both genes, we observed no fluorescence in uninjected embryos (Fig. 3F, I)

### Combinatorial expression of tissue-specific mNG2_1-10_ and mNG2_11_-tagged proteins

After successfully generating mNG2_1-10_ transgenic lines and mNG2_11_ insertions, we next determined whether these lines could be combined to achieve tissue-specific protein labeling (Fig. 4A). We crossed each of our mNG2_11_-tagged lines — *mNG2_11_-tubb4b* and *krt8-mNG2_11_* — with each of our mNG2_1-10_ transgenic lines — *fezf2:mNG2_1-10_*, *myl7:mNG2_1-10_*, and *ubb:mNG2_1-10_*. For *mNG2_11_-tubb4b*, crossing to *ubb:mNG2_1-10_* produced fluorescence broadly throughout the head (Fig. 4B), similar to mNG2_1-10_ mRNA injection and to the reported expression pattern for *tubb4b* (Thisse and Thisse, 2008; Zhuo et al., 2012). In contrast, crossing to *fezf2:mNG2_1-10_* resulted in fluorescence restricted to the brain and eye (Fig. 4C), consistent with the known expression pattern for *fezf2* (Jeong et al., 2006). Finally, crossing to *myl7:mNG2_1-10_* resulted in no observable fluorescence (Fig. 4D). This result is consistent with the reported expression pattern for *tubb4b*, which has not been reported to be expressed in the heart.

**Figure 4.**
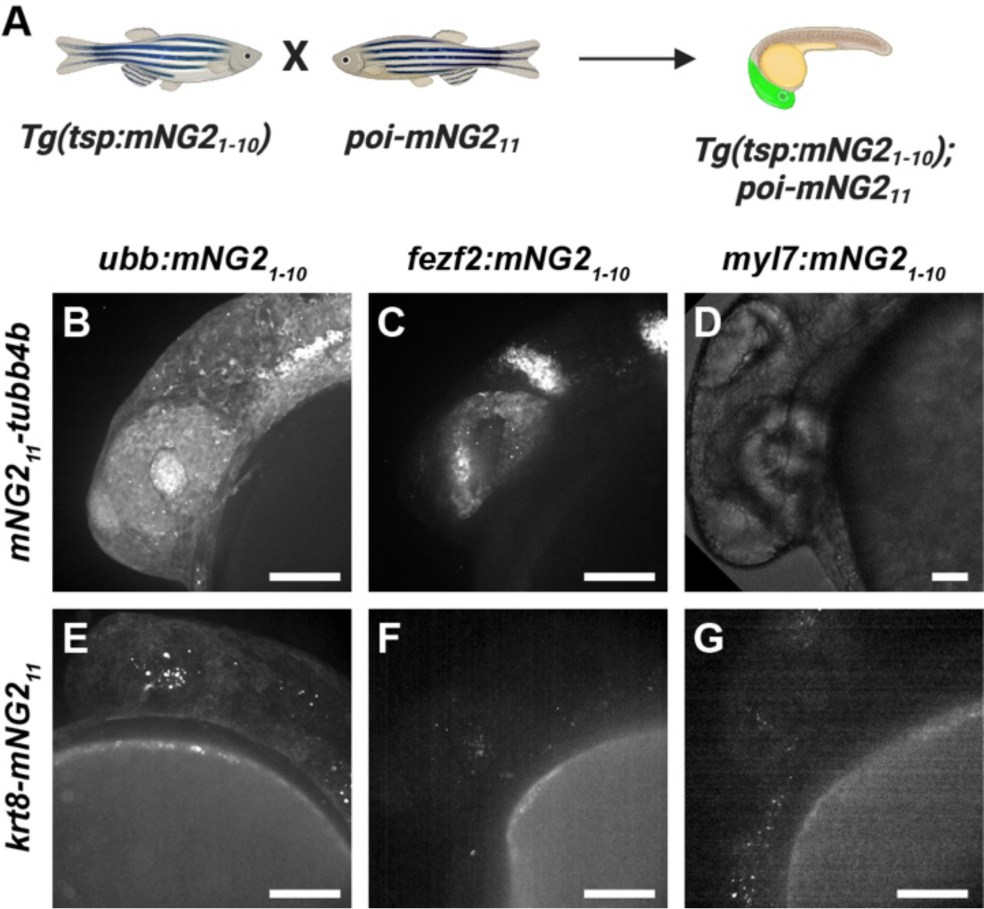
Combinatorial expression of tissue-specific mNG21-10 and mNG211-tagged proteins. **A.** Schematic of crossing strategy. Tsp, tissue-specific promoter. poi, protein of interest. **B-G.** Representative images of embryos resulting from crosses between *mNG211-tubb4b* (B–D) or *krt8-mNG211* (E–G) and *ubb:mNG21-10* (B, E), *fez2f:mNG21-10* (C, F), or *myl7:mNG21-10* (D, G). Images were acquired at 24 hours post-fertilization. Image in D has been overexposed to emphasize lack of fluorescence. Scale bars, 100 µm.

The results we obtained for *krt8-mNG2_11_* similarly demonstrated retention of endogenous expression patterns. Crossing to *ubb:mNG2_1-10_* resulted in fluorescence primarily in the skin (Fig. 4E), similar to mNG2_1-10_ mRNA injection and to the reported expression pattern for *krt8* (Fischer et al., 2014; Thisse and Thisse, 2008). And, crossing to both *fezf2:mNG2_1-10_* or *myl7:mNG2_1-10_* resulted in no observable fluorescence (Fig. 4F–G), which is expected as *krt8* has not been reported to be expressed in either cardiac or neural tissues.

Altogether, our results show that combining transgenic mNG2_1-10_ expression and mNG2_11_ tagging can achieve tissue-specific fluorescent protein labeling that preserves endogenous expression patterns.

### Directing protein localization with split-mNG2

Given that split-mNG2 fragments self-assemble, it may be possible to use mNG2_1-10_ as a “bait” to direct mNG2_11_-tagged proteins to specific subcellular locations. To determine the feasibility of this application, we fused mNG2_1-10_ to a localization signal for the outer mitochondrial membrane (mito-mNG2_1-10_) (Bear et al., 2000) (Fig. 5A). We then injected mRNA for mito-mNG2_1-10_ into *krt8-mNG2_11_* embryos. Compared to control embryos injected with untagged mNG2_1-10_ (Fig. 5B–D), embryos injected with mito-mNG2_1-10_ exhibited qualitatively different fluorescence localization patterns that co-localized with the mitochondrial dye MitoTracker (Fig. 5E–F). These results suggested that mito-mNG2_1-10_ was indeed directing mNG2_11_-tagged Keratin 8 to the mitochondria. Interestingly, *krt8-mNG2_11_* embryos injected with mito-mNG2_1-10_ exhibited a higher frequency of defects consistent with mislocalization of Keratin 8 protein including skin blistering (Fig. 5I) and collapsed fin folds (Fig. 5J). These defects were observed in 17.3% of embryos injected with mito-mNG2_1-10_ (n=248 embryos) compared with 1.1% of uninjected embryos (n=90 embryos). Together, these results demonstrate that by anchoring mNG2_1-10_ to specific cellular compartments, the split-mNG2 system can be used to direct protein localization and perturb protein function.

**Figure 5.**
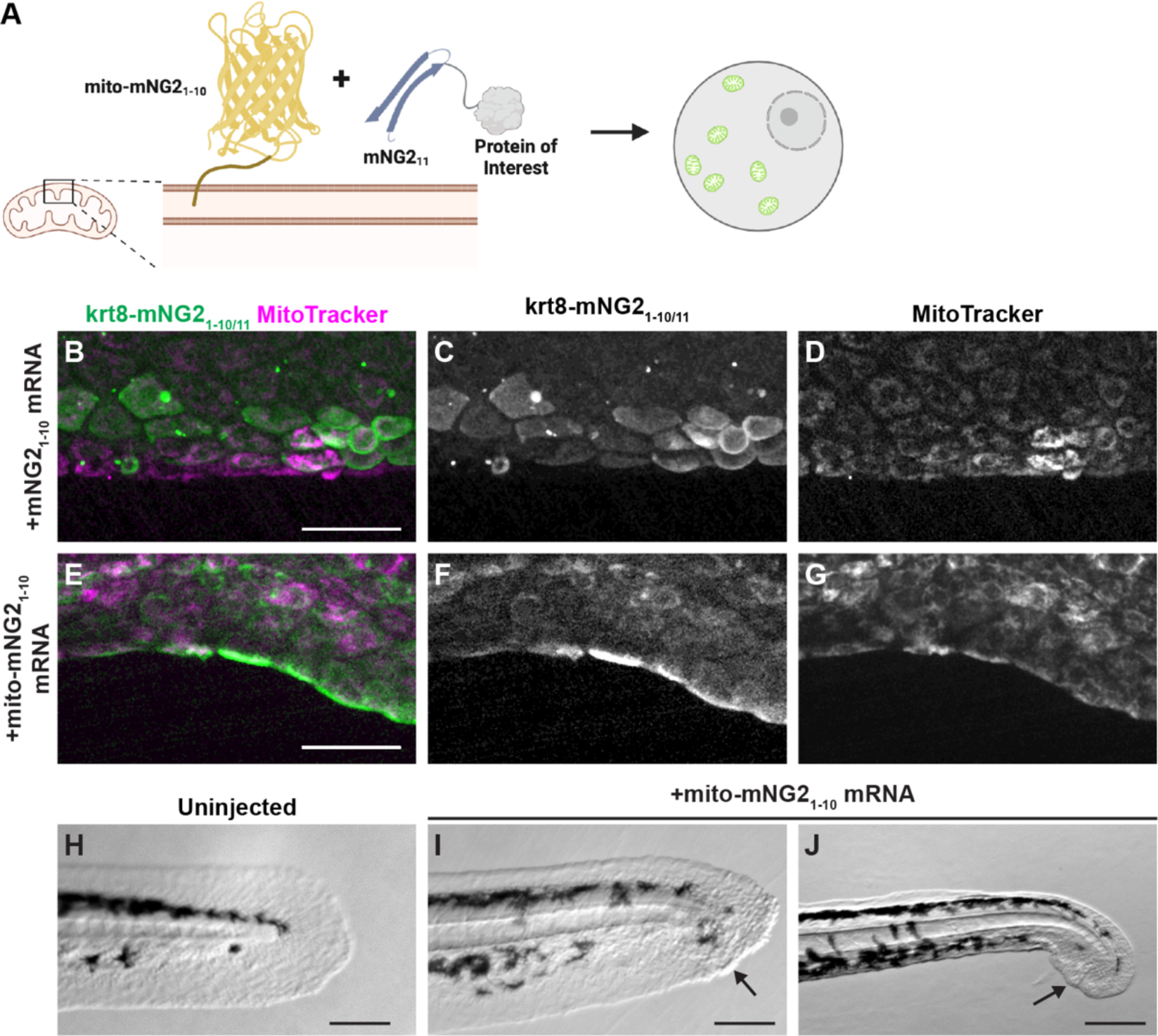
Directing protein localization with split-mNG2. **A.** Schematic illustrating use of the split-mNG2 system to sequester proteins of interest on mitochondria. **B–G.** Representative images of *krt8-mNG211* embryos injected with mNG21-10 (B–D) or mito-mNG21-10 (E–G) mRNA. Images were acquired from the tail fin epidermis at 48 hours post-fertilization (hpf). Mitochondria were labeled with MitoTracker dye. Scale bars, 50 µm. **H–J.** Representative images of the tails of uninjected control embryos (H) *krt8-mNG211* embryos injected with mito-mNG21-10 mRNA (I, J). Images were acquired at 48 hpf. Arrows denote blisters. Scale bars, 200 µm.

## Discussion

In this study, we describe using the mNG2_1-10/11_ split fluorescent protein system to achieve tissue-specific fluorescent labeling of endogenous proteins in zebrafish embryos. We further demonstrate that the split-mNG2 system can be used to control protein localization by anchoring the mNG21-10 fragment to specific cellular compartments.

Similar FP_1-10/11_ systems are now commonly used as endogenous protein labeling tools in cell lines (Cho et al., 2022; Feng et al., 2017; Kamiyama et al., 2016; Leonetti et al., 2016). The popularity of these systems is primarily due to the ease with which the short FP_11_ sequences can be inserted into gene loci. The general utility of split-FP systems for protein labeling has also been demonstrated in multicellular organisms including zebrafish (Kesavan et al., 2021) and mouse embryos (O’Hagan et al., 2021), but in these studies the corresponding FP_1-10_ fragment was delivered constitutively. Tissue specificity has been achieved in *C. elegans* (Goudeau et al., 2021; He et al., 2019; Hefel and Smolikove, 2019; Noma et al., 2017) and *Drosophila* (Kamiyama et al., 2021) and now in zebrafish (this study).

Control of protein localization is a novel application of the split-mNG2 system. As we demonstrate by tethering Keratin 8 to mitochondria, this approach can be used for manipulating protein function by sequestering proteins away from their normal site of function. But the same approach could also be used to force constitutive localization of a protein to its site of activity or to direct ectopic localization. Reconstituted fluorescent proteins tagged with specific localization sequences retain their fluorescence, thus, successful mislocalization can easily be confirmed. When combined with transgenic expression of mNG2_1-10_, these perturbations can be applied to specific tissues of interest for even broadly expressed proteins.

Previous reports have suggested that not all proteins can be easily labeled with the split-mNG2 system (Cho et al., 2022; Leonetti et al., 2016; O’Hagan et al., 2021). Fluorescent labeling may fail because the target protein does not tolerate mNG2_11_ tagging. Thus, mNG2_11_ fusion proteins should be designed using the same considerations as with any epitope tag.

Even if tagging is tolerated, some proteins may not be expressed at high enough levels to produce a detectable fluorescent signal (Leonetti et al., 2016; O’Hagan et al., 2021). This challenge could be overcome by inserting multiple repeats of the mNG2_11_ sequence to increase fluorescent signal as has been demonstrated for split-GFP (He et al., 2019; Hefel and Smolikove, 2019; Kamiyama et al., 2016, 2021; Noma et al., 2017). Additionally, a third generation split-mNG system was recently developed and reported to have improved spectral properties (Zhou et al., 2020), which may extend the use of split-FP labeling to low or moderately expressed proteins.

In this study, we focused on the use of split-mNG2 as a protein labeling tool. However, the ability to control expression of of these protein fragments independently, paired with their ability to self-assemble, could be leveraged for other applications. For example, they could be used as coincidence detectors to monitor cell states or signaling pathway activation. And because fluorescence is only produced when the two fragments bind, they could be used to visualize interactions at multiple length scales, i.e., between proteins, organelles, cells, or adjacent tissues.

In summary, we have demonstrated that the split-mNG2 system can function in zebrafish to endogenously label proteins in a tissue-specific manner, with other potential applications that make it broadly useful to many areas of investigation.

## Materials and Methods

### Zebrafish strains

Adult *Danio rerio* zebrafish were maintained under standard laboratory conditions. Zebrafish in an outbred *AB*, TL, or *EKW* background were used as wild-type strains. Strains generated in this study are: *Tg(fezf2:mNG2_1-10_)^ucm120^*; *Tg(myl7:mNG2_1-10_)^ucm121^*; *Tg(ubb:mNG2_1-10_)^ucm117^*; *krt8^ucm132^*; and *tubb4b^ucm132^*. This study was performed with the approval of the Institutional Animal Care and Use Committee (IACUC) of the University of California Merced (Protocol #2023-1144).

### mRNA expression

All expression plasmids for *in vitro* mRNA synthesis were generated in a pCS2 backbone. To generate pCS2-GFP_1-10_, GFP_1-10_ was PCR amplified from pACUH-GFP_1-10_ (Bo Huang, University of California San Francisco) and cloned into pCS2 by enzymatic assembly (Gibson et al., 2009). To generate pCS2-mNG2_1-10_, mNG2_1-10_ was PCR amplified from pSFFV-mNG2_1-10_ (Bo Huang, University of California San Francisco) and cloned into pCS2 by enzymatic assembly. To generate pCS2-GFP_11_-H2B and pCS2-mNG2_11_-H2B, GFP_11_ (5ʹ-CGTGACCACATGGTCCTTCATGAGTATGTAAATGCTGCTGGGATTACA-3ʹ) and mNG2_11_ (5ʹ-ACCGAGCTCAACTTCAAGGAGTGGCAAAAGGCCTTTACCGATATGATG-3ʹ) were directly synthesized by Integrated DNA Technologies and H2B was PCR amplified from GFP-H2B (Hesselson et al., 2009); fragments were fused and cloned into pCS2 by enzymatic assembly. To generate pCS2-mito-mNG2_1-10_, the outer mitochondrial membrane signal sequence was PCR amplified from pMSCV-FPPPP-mito (Bear et al., 2000) and cloned into pCS2-mNG2_1-10_ by enzymatic assembly. Capped messenger RNA was synthesized using the mMESSAGE mMACHINE kit (Ambion), and 500 pg of each mRNA was injected at the one- or two-cell stage.

Generation of mNG2_1-10_ transgenic lines

All transgene plasmids were generated in a pµTol2 backbone (LaBelle et al., 2021). mNG2_1-10_ and promoter sequences for *fezf2* (Berberoglu et al., 2009), *myl7* (Huang et al., 2003), or *ubb* (Mosimann et al., 2011) were PCR amplified then fused and cloned into pµTol2 by enzymatic assembly to generate pµTol2-fez:mNG2_1-10_, pµTol2-myl7:mNG2_1-10_, and pµTol2-ubb:mNG2_1-10_, respectively. The constructs were used to generate *Tg(fez:mNG2_1-10_)^ucm120^*; *Tg(myl7:mNG2_1-10_)^ucm121^*; *Tg(ubb:mNG2_1-10_)^ucm117^* using standard transgenesis protocols (Clark et al., 2011; Kawakami, 2004).

### CRISPR/Cas-directed insertion of mNG2_11_

Guide RNAs (gRNAs) were designed using CRISPRscan (Moreno-Mateos et al., 2015) and synthesized as previously described (Varshney et al., 2016). The double-stranded DNA template for homology directed repair was assembled from two oligomers synthesized by Integrated DNA Technologies. Each oligomer contained the sequence for mNG2_11_, a 10-amino acids-encoding linker sequence (5’-GGAGCTGGTGCAGGCGCTGGAGCCGGTGCC-3’), and a homology arm. Oligomers were hybridized to obtain a double-stranded template with single-stranded, 30 bp-long homology arms at each end (Liang et al., 2017). gRNAs, donor DNA, and Cas9 mRNA were injected at the one-cell stage as previously described (Gagnon et al., 2014).

To verify insertion, we pooled 40 injected embryos at 24 hpf, isolated genomic DNA, and performed PCR using two sets of primer pairs per gene covering the 5’ and 3’ insertion sites.

The same primer sets were used for quantitative PCR (qPCR) to estimate knock-in efficiency. Each qPCR reaction contained 2X PerfeCTa® SYBR Green FastMix (Quantabio), five-fold diluted genomic DNA, and 325 nM of each primer. Reactions were carried out on a QuantStudio3 (Applied Biosystems) real-time PCR machine using the following program: initial activation at 95°C for 10 min, followed by 40 cycles of 30 s at 95°C, 30 s at 60°C, and 1 min at 72°C. Once the PCR was completed, a melt curve analysis was performed to determine reaction specificity. The gene *prox1a* was used as a reference. Primers used in this study (presented 5’–3’):

5’ h2az2b-mNG211 forward: TTGTGTGTTTGTGCGTCCGC

5’ h2az2b-mNG211 reverse: GCCACTCCTTGAAGTTGAGC

3’ h2az2b-mNG211 forward: GCTCAACTTCAAGGAGTGGC

3’ h2az2b-mNG211 reverse: ACGAAGCCCCGAAAGCACAC

5’ mNG211-krt8 forward: ATACAGCGGCGGATACAGCG

5’ mNG211-krt8 reverse: GCCACTCCTTGAAGTTGAGC

3’ mNG211-krt8 forward: GCTCAACTTCAAGGAGTGGC

3’ mNG211-krt8 reverse: AAGGCACGACAAGAGCGGTG

5’ mNG211-tubb4b forward: CACATCTCGAATTACGACCTCA

5’ mNG211-tubb4b reverse: GCCTTTTGCCACTCCTTGAAG

3’ mNG211-tubb4b forward: GCTCAACTTCAAGGAGTGGC

3’ mNG211-tubb4b reverse: AAAACAAGCAAGGATTAGCGTC

prox1a forward: TGTCATTTGCGCTCGCGCTG

prox1a reverse: ACCGCAACCCGAAGACAGTG

To verify germline transmission and establish stable lines, injected F_0_ embryos were raised to adulthood then outcrossed to wild-type zebrafish. We pooled 40 of the resulting F_1_ embryos at 24 hpf, isolated genomic DNA, and performed PCR using the same primer sets as above. PCR fragments were cloned in pGEM-T (Promega), and the inserts were sequenced by Sanger sequencing (University of California Berkeley DNA Sequencing Facility). Only clutches containing precise insertion of the mNG2_11_ plus linker sequence were kept for propagation. At adulthood, individual F_1_ zebrafish were genotyped by fin clipping using the same primer sets as described above. Only animals containing precise insertion of the mNG2_11_ sequence were kept for line propagation.

### Microscopy and image processing

Fluorescence and brightfield images were acquired on an Olympus SZ51 stereomicroscope equipped with a DP23 monochrome camera and cellSens software (Evident) or with an Olympus IX83 microscope equipped with a 10x/0.4NA or 30x/1.05 NA objective (Evident), a spinning-disk confocal unit (CSU-W1; Andor), a scientific complementary metal– oxide–semiconductor (sCMOS) camera (Prime 95b; Teledyne Photometrics), and MicroManager software (Edelstein et al., 2014). Dechorionated embryos or larvae were embedded in 1.5% low-melting agarose (ISC BioExpress) containing 0.01% tricaine (Sigma-Aldrich) within glass-bottom Petri dishes (MatTek Corporation). For mitochondria labeling, embryos were incubated in 50 nM MitoTracker Red CMXRos (Invitrogen) for 30 minutes prior to agarose embedding. Standard filter settings were applied and brightfield and fluorescence images were merged after acquisition. Identical exposure settings for fluorescence images were used for all embryos from the same set of experiments. Images were processed in Fiji (Schindelin et al., 2012) as follows: denoised using the Non-local Means Denoise plugin (Buades et al., 2005), maximum intensity Z-projection, brightness and contrast levels adjusted, converted to 8-bit depth, and cropped. Illustrations were created with BioRender (https://biorender.com).

## Acknowledgements

We thank Bo Huang (University of California San Francisco) for providing the split-GFP and split-mNG2 plasmids and advice, Anne Pipathsouk (University of California San Francisco) and Manuel Leonetti (Chan Zuckerberg Biohub) for helpful discussions, the Department of Animal Research Services (University of California Merced) for excellent fish care, and members of the Woo and Materna labs for helpful comments on the manuscript. This work was supported by grants from the Society for Developmental Biology, the National Institutes of Health (NIH R15HD102829), and the National Science Foundation (NSF-IOS-2238304) to S.W. J.L. was supported by NIH training grant T32GM141862. K.S. received support from the NSF-CREST: Center for Cellular and Biomolecular Machines at the University of California, Merced (NSF-HRD-1547848).

---

**Supplemental Figure 1.**
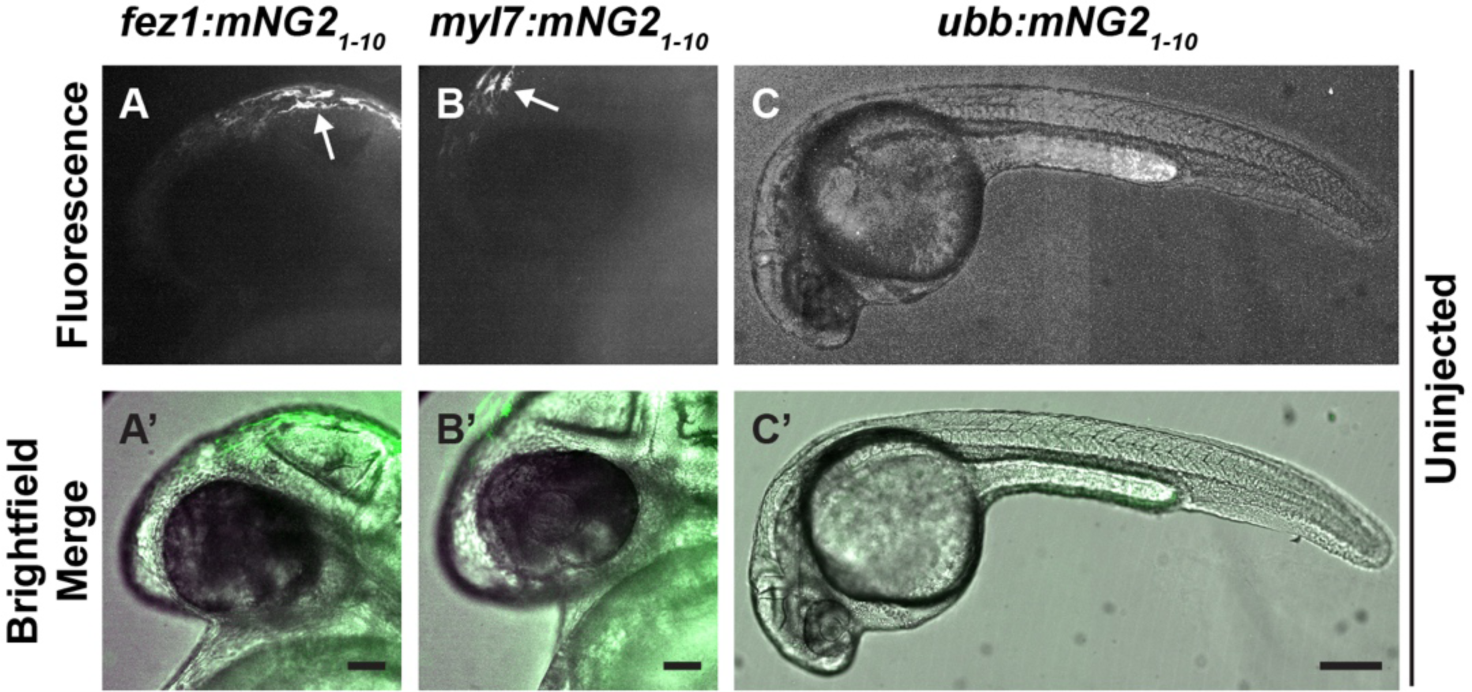
mNG21_-10_ transgenic lines are not fluorescent in the absence of mNG2_11_. **A–C.** Uninjected transgenic embryos expressing mNG2_1-10_ under control of the *fez1* (A–Aʹ), *myl7* (B–Bʹ), or *ubb* (C–Cʹ) promoters. Images in A, B, and C have been overexposed to emphasize absence of fluorescent signals other than autofluorescent pigment cells (arrows in A, B). Scale bars in Aʹ and Bʹ, 50 µm. Scale bar in Cʹ, 200 µm.

**Supplemental Figure 2.**
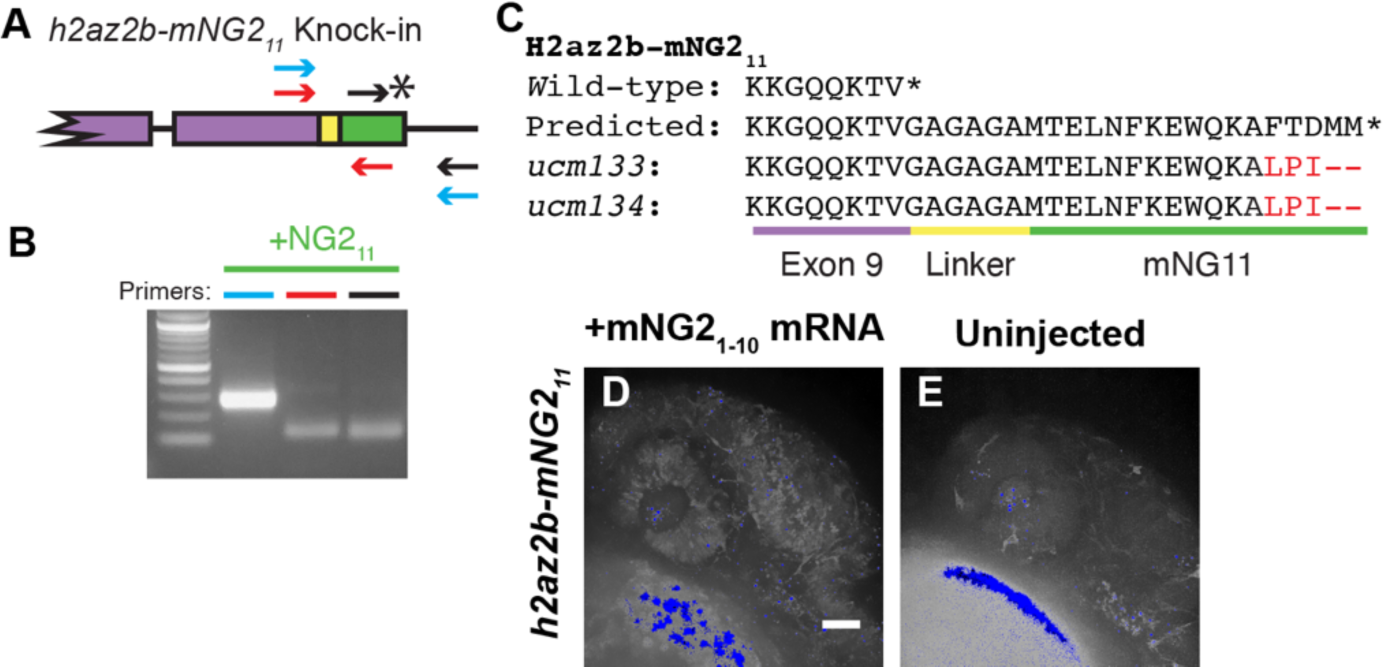
mNG211 tagging of *h2az2b*. **A.** Schematic of mNG2_11_ insertion into the *h2az2b* gene. Purple, endogenous exon sequence. Yellow, linker. Green, mNG2_11_. Asterisk, stop codon. Arrows denote primers used in B. **B.** mNG2_11_ insertion was assessed by PCR. The primers used correspond to the arrows shown in A. **C.** Amino acid sequences of wild-type, predicted mNG2_11_ fusion, and recovered alleles for H2az2b. Mismatches between the predicted and recovered sequences are highlighted in red. Asterisk, stop codon. **D–E.** Representative images of *h2az2b-mNG2_11_* embryos at 24 hours post-fertilization. Embryos injected with mNG2_1-10_ show dim nuclear-localized fluorescence (D) compared to uninjected embryos (E). Very bright spots (blue) are likely autofluorescent yolk and debris. Scale bar, 50 µm.

